# Time Cells in the Human Brain Support Working Memory Maintenance

**DOI:** 10.64898/2026.01.12.699074

**Authors:** Xiaoxuan Xiao, Ueli Rutishauser, Taufik A. Valiante, Jiannis Taxidis

**Affiliations:** Krembil Research Institute, University Health Network, Toronto, ON, Canada; Institute of Biomedical Engineering, University of Toronto, Toronto, ON, Canada; Program in Neurosciences and Mental Health, The Hospital for Sick Children, Toronto, ON, Canada; Department of Neurosurgery, Cedars-Sinai Medical Center, Los Angeles, CA, USA; Division of Biology and Biological Engineering, California Institute of Technology, Pasadena, CA, USA; Center for Neural Science and Medicine, Cedars-Sinai Medical Center, Los Angeles, CA, USA; Center for Advancing Neurotechnological Innovation to Application (CRANIA), Toronto, ON, Canada; Max Planck-University of Toronto Centre for Neural Science and Technology, Toronto, ON, Canada; Institute of Medical Sciences, University of Toronto, Toronto, ON, Canada; Division of Neurosurgery, Department of Surgery, University of Toronto, Toronto, ON, Canada; Department of Physiology, University of Toronto, Toronto, ON, Canada

**Keywords:** human single neurons, working memory, time cells, concept cells, time encoding

## Abstract

Working memory (WM), the active retention of information over short periods, is a fundamental cognitive function, yet its underlying neural mechanisms remain unclear. In rodents, cue-selective “time cells” fire at specific timepoints after a WM memory cue, collectively forming sequences that encode cue-memory and elapsed time, providing a temporal code for maintaining information across time intervals. Whether similar dynamics support WM in the human brain is unknown. Here, we analyzed intracranial single-neuron recordings from medial frontal and medial temporal regions of patients performing a WM task. We found time cells with temporally tuned activation during WM maintenance, collectively forming robust sequences that tiled a delay period. Time-cell coordination in the hippocampus predicted successful WM maintenance, whereas in the pre-supplementary motor area it reflected memory load. Furthermore, we identified distributed cue-selective time cells that encoded both the identity of a memorandum and elapsed maintenance time, providing a temporally structured mnemonic code that complements persistent firing of concept cells. Together, these findings establish time-cell sequences as a conserved neural mechanism supporting human working memory.

## Introduction

Working memory (WM), the transient and active retention of information in memory, is central to cognitive flexibility and planning^1–3^. Despite decades of research, the neural mechanisms that sustain WM representations remain little understood, particularly in the human brain.

Classic models propose that WM is supported by persistent, stimulus-specific firing of single neurons, most prominently in prefrontal and medial temporal cortices in humans and non-human primates^4–6^. However, growing experimental and theoretical work suggests that persistent firing alone may not fully account for WM maintenance. Instead, WM representations can be embedded within dynamic population activity, including ramping responses and sequentially activated neural ensembles^7–18^.

Among these dynamic mechanisms, “time cells” have emerged as a compelling candidate for organizing memory across delays. Time cells have been observed in the rodent brain, across cortical and subcortical regions, predominantly hippocampus and entorhinal cortex^7,8,19–22^. Each time cell fires reliably at a circumscribed timepoint within a delay interval, and together these cells form reproducible sequences tiling the delay interval and encoding elapsed time on the scale of seconds^22,23^. These sequences provide a temporal scaffold for organizing experiences and can bridge discontiguous events in memory^7,8,23,24^.

Importantly, time cells can also be memoranda-specific, i.e. only fire after a particular cue (‘cue’ referring to the identity of a memorandum throughout), thereby encoding both memory of the cue and time since its presentation^17^. Such cue-specific sequences emerge during learning of a WM task^17^ and reflect expected delay durations^17,18^, providing a mechanism for binding mnemonic content to temporal context during WM maintenance^21,25,26^.

Despite extensive evidence in rodents, it remains unknown whether analogous time-cell sequences exist in the human brain and, if so, whether they contribute to WM. Prior human single-neuron studies have reported temporally modulated firing during navigation^27^ and episodic memory tasks^28^, but these paradigms required explicit temporal structure and did not isolate WM maintenance in the absence of sensory input. As a result, it remains unclear whether the human brain can internally generate time-cell sequences to support WM.

Here, we address this gap by analyzing intracranial single-neuron recordings from medial frontal and temporal regions of neurosurgical patients performing a WM task with images as memoranda^6,29^. We identify robust time-cell sequences that tile the WM maintenance interval, despite the absence of any timing requirement by the task. We show that hippocampal time-cell activity predicts successful WM maintenance, whereas time-cell dynamics in pre-supplementary motor area reflect memory load and subsequent retrieval speed to a memory probe question.

Furthermore, we identify cue-selective time cells that encode both memorandum identity and elapsed time, providing a temporally structured mnemonic code that complements persistent firing of classical concept cells^6^.

Together, our findings establish time-cell sequences as a core mechanism of human WM and demonstrate a conserved temporal coding principle that links rodent and human memory systems.

## Results

### Time Cells During WM Maintenance

To investigate whether time cells exist in the human brain during WM, we analyzed single-neuron recordings from the Kyzar et al. (2024) open-source dataset^29^. This dataset contains spike times from neurons in the medial temporal lobe and medial frontal lobe of 21 neurological patients, during performance of a WM task.

Participants were shown 1–3 images sequentially (encoding period; Load 1-3 respectively), followed by a variable 2.5–2.8s long delay period (maintenance period), and then a probe image. The task was to report if the probe was identical to one of the images shown during the encoding period of the same trial (Fig. 1A). Subjects performed well (88.7% ± 5.7% accuracy; mean ± SD), with reaction times increasing with memory load^29^ as expected for a WM-dependent behavior (Fig. 1B). Single neuron recordings (n = 902 units) were obtained from depth electrodes implanted in five brain regions involved in memory and cognitive control: hippocampus, amygdala, pre-supplementary motor area (PSMA), dorsal anterior cingulate cortex (dACC), and ventromedial prefrontal cortex (vmPFC; Fig. 1A).

**Fig. 1.**
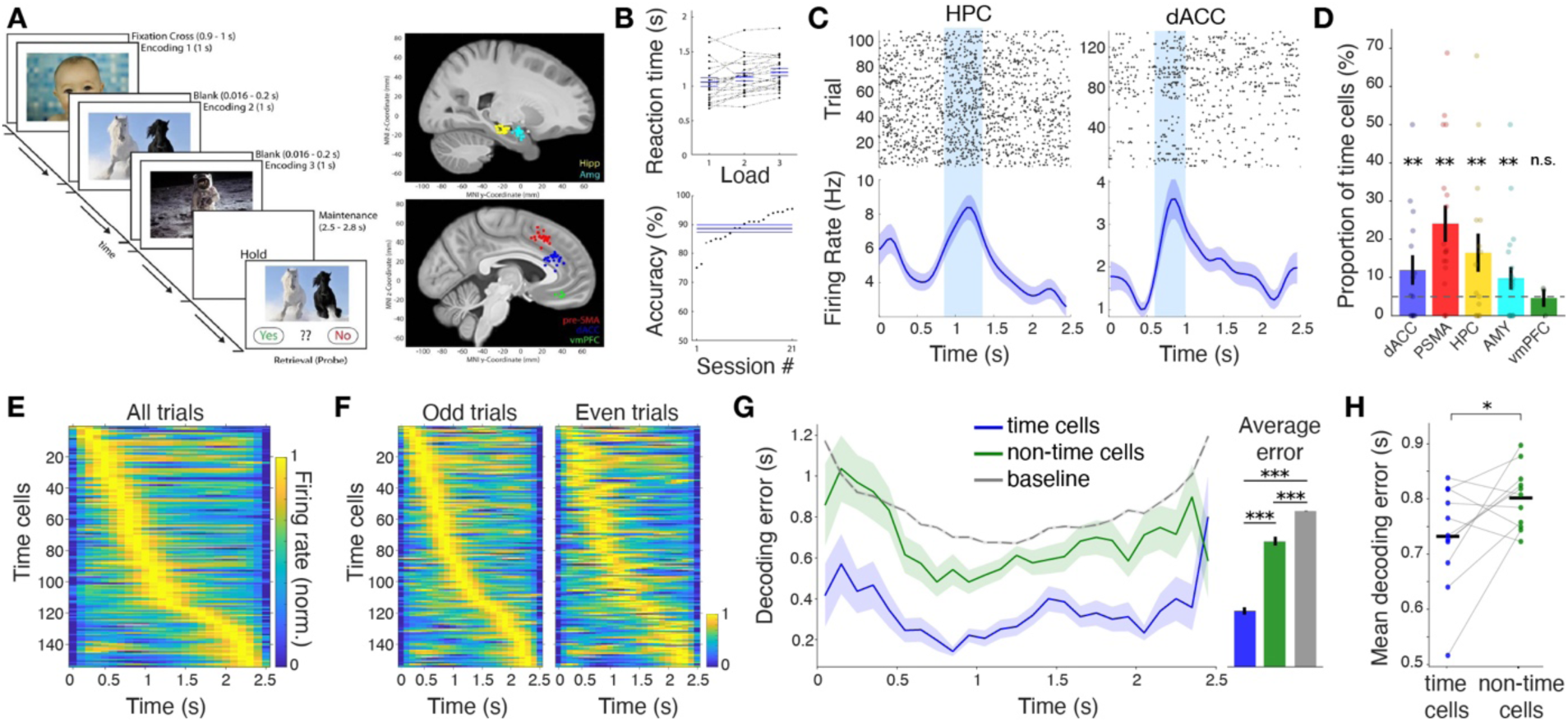
| Time-cells in the human brain during WM maintenance. **A.** Sternberg task: After a fixation baseline, 1–3 images (Load 1-3 respectively) were shown sequentially (1 s, 0.5 s inter-stimulus), followed by a variable maintenance delay (2.5-2.8 s) and then a probe image. Participants reported whether the probe matched any of the samples. Right: Stereotactic locations of microwire bundles (dots) plotted in MNI152 space and overlaid on the CIT168 atlas. Colours denote brain region: dorsal anterior cingulate cortex (dACC, blue), pre-supplementary motor area (pre-SMA, red), hippocampus (HPC, yellow), amygdala (AMY, cyan), and ventromedial prefrontal cortex (vmPFC, green). For visualization, all sites are projected to the right hemisphere. **B.** Behaviour. Top: Median reaction time for each load condition, relative to the probe image onset. Dashed lines link individual participants. Bottom: Task performance across participants, rank ordered. **C.** Raster plots (top) and mean ± s.e.m. firing rate (bottom) across the 2.5s long maintenance period for two example time cells. Hereafter, t = 0 denotes maintenance period onset. Blue bands indicate their time field ± 0.5 SD above baseline. **D.** Mean ± s.e.m proportion of time-cells per brain region in each patient (dots). Dashed line: Shuffle baseline. Empirical p_dACC_ = 0.001996, p_PSMA_ = 0.001996, p_HPC_= 0.001996, p_AMY_ = 0.001996, p_vmPFD_ = 0.44511. **E.** Trial-averaged normalized firing-rates for all time-cells (rows), stacked by time field, across the maintenance phase. **F.** Same for firing rates across odd and even trials only, sorted by their peak timepoint in odd trials. **G.** Mean ± s.e.m absolute error in Bayesian decoding of maintenance time, using time-cells (blue), all other neurons (green) and shuffle baseline (grey; see Methods). Right: corresponding mean error across time bins and trials (time tells vs non-time cells: t = -12.231, p = 6.153 x10^-33^; time cells vs shuffle baseline: t = -27.725, p = 4.20910^-120^; non-time cells vs shuffle baseline: p = 1.29810^-4^; two-tailed t-test:). **H.** Mean absolute error in Bayesian decoding of maintenance time, using time-cells vs non-time cells, per patient (lines; patients with ≥5 time cells included; t = -2.318, p = 0.043, paired two-sided t-test). Panels **A-B** are reproduced^1^ with permission from the corresponding author.

We identified time cells as neurons with significant firing rate increases across all trials at specific timepoints (time field) within the 2.5 sec long maintenance interval (Fig. 1C; see Methods). In total, 152/902 neurons met this criterion. Time cells were observed at numbers larger than expected by chance in all recorded regions except vmPFC, which did not contain time cells exceeding chance levels (2/32 units; p > 0.05 versus shuffled baseline; see Methods; Fig. 1D). The highest proportion of time cells was observed in PSMA (66/250 units), followed by hippocampus (31/190), dACC (27/171) and amygdala (28/259). Across patients, the mean proportion of time cells was 16.1% ± 2.7% (mean ± SEM).

Pooled together, time cells formed sequences of temporally modulated activity that tiled the entire maintenance period (Fig. 1E). To validate the stability of time cell activation across trials, we sorted time cells by their peak activation in odd trials only and confirmed that they maintained similar peak firing times in even trials (Fig. 1F).

To further test the robustness of the encoding of time, we trained a Bayesian decoder to predict time elapsed since start of the maintenance period from neural firing rates of different groups of cells. Throughout the maintenance period, time cells yielded significantly lower errors in decoded time compared to non-time cells or to shuffle baseline; both across pooled trials and time bins (Fig. 1G) and on a per-patient level (Fig. 1H).

To test whether time cell sequences are specific to time intervals of WM maintenance or also exist in other types of memory tasks, we repeated our analysis in a separate public dataset involving episodic memory encoding^30^. Participants viewed short video clips which contained predefined “hard” boundaries (‘HB’; transitions between unrelated scenes), “soft” boundaries (‘SB’; camera shifts within a scene) or no boundaries (‘NB’; Fig. S1), followed by a memory test on the content of the viewed clips. Single-unit recordings were obtained from the hippocampus (n = 343 neurons), amygdala (n = 169 neurons), and parahippocampal gyrus (n = 68 neurons) during the task. Using the same time cell detection method as before, applied across the first 4 sec of HB and NB videos, cells in all three regions were detected that had a time field: hippocampus (42/343, 12.20%), amygdala (21/169, 12.40%), and parahippocampal gyrus (5/68, 7.35%) (Fig. S1). However, these cells either encoded video initiation (time fields at < 1sec) or exhibited persistent-like firing throughout the 4 sec from clip onset, resulting in chance-level time decoding 1 sec after clip onset (Fig. S1). Moreover, the same sequence did not repeat after a HB event, confirming these early temporally modulated cells encoded video onset, instead of time from any visual boundary. When detecting time cells throughout the entire HB videos, temporally modulated neurons also appeared within 1 sec after the boundary onset, corresponding to the previously described ‘boundary cells’^30^, but were not followed by a time cell sequence, unlike in the WM task (Fig. S1).

Collectively, our findings reveal distributed time cells in the human brain, selectively during WM maintenance, carrying temporal information which may be relevant to memory retention.

### Time Cell Activation Correlates with WM Performance

Persistent spiking of medial temporal neurons during the maintenance period was previously found to correlate with WM performance^6^. We thus tested whether time cell activation differed between trials with successful versus unsuccessful WM maintenance as assessed by the accuracy of the answer to the probe question.

Time cells often exhibited more strongly modulated firing rates in correct trials compared to incorrect ones (trials with incorrect probe responses; Fig. 2A). Indeed, the mean z-scored firing rate within 100ms around each cell’s field, was significantly higher in correct than incorrect trials (Fig. 2B). When repeating this comparison in each brain region, we found that, on average, this effect was significant only in dACC and hippocampal time cells (Fig. 2C), suggesting stronger modulation of these cells with successful memory maintenance. Overall, this resulted in more pronounced spiking sequences in correct than incorrect trials (Fig. 2D).

**Fig. 2.**
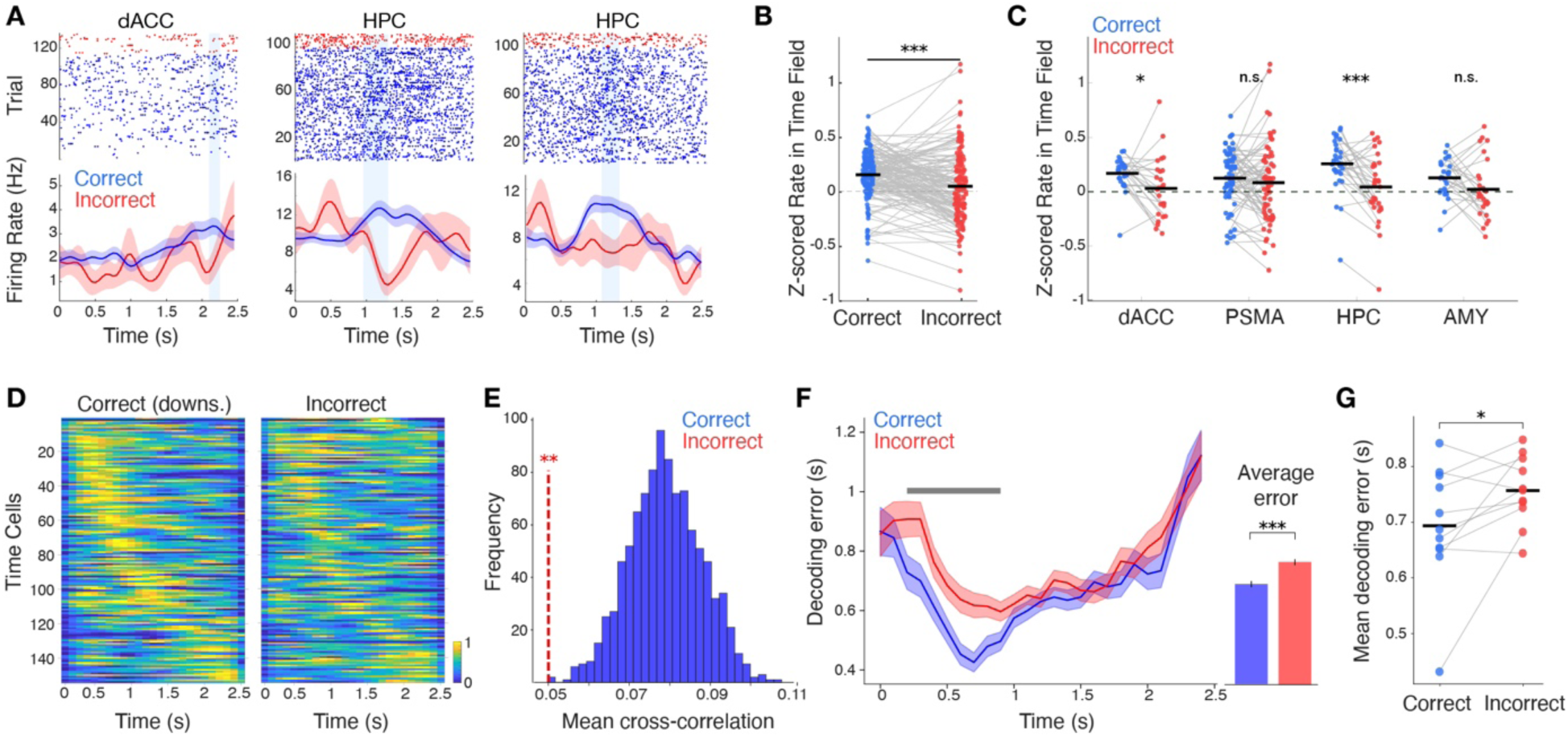
| Time cell activation predicts WM performance. **A.** Raster plot and mean± SEM firing rates of example time cells across the maintenance period in correct (blue) and incorrect trials (red). **B.** Average z-scored firing rates within each cell’s time field during correct vs incorrect trials (t = 4.25, p value = 3.6459 * 10^-5^, two-tailed paired-sample t-test). **C.** Same as in **B** but for time cells grouped by brain region. (p_dACC_ = 0.0207; p_PSMA_ = 0.293; p_HPC_ = 0.00031; p_AMY_ = 0.0569; two-tailed paired-sample t-tests). **D.** Average normalized firing rates of time cells during correct (d) and incorrect (e) trials, sorted by time field across trials. Correct trials were randomly downsampled to match incorrect ones. **E.** Mean pairwise cross-correlations of time-cell z-scored firing rates in incorrect trials (red) vs distribution of cross-correlations in correct trials (size-matched random subsampling; n = 1000 iterations; empirical p value = 0.001; see Methods). **F.** Mean ± s.e.m. absolute error in time decoding by Bayesian decoders trained on correct-trial data and tested on held-out correct (blue) and incorrect (red) trials. Grey bars indicate time points where correct and incorrect trials differ at p < 0.05. Right: Corresponding mean decoding error across time bins and trials. (t = -5.12, p = 3.080e-07; two-sided t-test). **G.** Mean decoding error per patient (lines; t = -2.34, p = 0.041; paired-sample two-sided t-test).

To quantify the temporal coordination across time cell sequences, we computed average cross-correlations of z-scored firing rates between all pairs of simultaneously recorded time cells within individual trials, at time lag equal to the distance between their time fields. To account for the low sample of incorrect trials, this was done in random size-matched subsets of correct trials, with resampling. The mean cross-correlation of time cells was significantly higher in correct than incorrect trials (p < 0.01, Fig. 2E).

To further assess whether the encoding of temporal information on the population level is correlated with successful WM, we trained a Bayesian decoder to decode time elapsed since the start of the maintenance period, based on the firing of time cells in a subset of correct trials. The classifier was then tested on incorrect trials and the remaining correct ones (sample sizes matched). Time decoding errors were significantly lower in correct than incorrect trials throughout the earlier maintenance period (Fig. 2F), resulting in lower mean errors across pooled time bins (Fig. 2F) and at per-patient level (Fig. 2G). To assess this effect per-brain region (while accounting for low numbers of time cells per patient in most regions), we compared time decoding errors from Bayesian decoders trained on pooled time cells from all regions versus after removing time cells from a region (see Methods). Removing hippocampal time cells, led to significantly lower improvement in decoding accuracy in correct versus incorrect trials, whereas removing time cells from other regions did not yield significant changes (Fig. S2), confirming that hippocampal time cells exhibited the strongest modulation by successful WM.

Moreover, to exclude any effects by memory load, we resampled correct trials to match the load distribution of incorrect trials. This did not alter our previous findings on z-scored firing rates within-field (p_all_ = 0.0002, p_dACC_ = 0.029, p_HPC_ = 0.0003), on cross-correlation analysis (empirical p = 0.001), on removing hippocampal time cells from Bayesian time-decoding (p = 0.0354), and on pooled decoding results (p = 7.759 x 10^-4^).

We then asked, for correct trials, whether time-cell modulation was related to the time it took participants to answer the probe question in a given trial. This reaction time reflects the efficiency of the WM representation, making it a sensitive metric of WM quality^31,32^. To do so, we compared time cell modulation between the fastest 30% and slowest 30% of correct trials in each load, pooled across loads (Figure S3). Time cells exhibited reduced temporal modulation during slow trials (Fig. S3). On average, z-scored within-field firing was significantly higher in fast than in slow correct trials, with the strongest per-region effects observed in PSMA and amygdala (Fig. S3). Consistent with this, a support vector machine (SVM) classifier trained on within-field firing rates of pooled time cells reliably discriminated fast from slow trials (Fig. S3), demonstrating that coordinated temporal tuning during maintenance is predictive of WM quality.

Collectively, our findings indicate that temporal coding, predominantly in hippocampal units, reflects successful WM maintenance, whereas in PSMA and amygdala reflect quality of memory maintenance, measured by reaction time.

### Time Cell Activation in PSMA is Mediated by Memory Load

Persistent spiking of medial frontal neurons during the maintenance period was shown to correlate with memory load^6^. We thus next examined whether time-cell firing patterns also varied between different load conditions.

Time cells often displayed sharply tuned activation during Load 1 and 2 trials but flattened or absent peaks during Load 3, particularly in early-maintenance time fields (Fig. 3A), suggesting a disruption in temporal coding in Load 3 trials. To quantify this disruption, we compared z-scored firing rates within each neuron’s time field across load conditions. On average, time cells in PSMA exhibited significantly lower temporal modulation during Load 3 trials, compared to Loads 1 and 2 trials, whereas time cells in other brain regions were not significantly affected by load (Fig. 3B). As a result, PSMA time-cell sequences were degraded in Load 3 trials, most notably among early time fields (Fig. 3C).

**Fig. 3.**
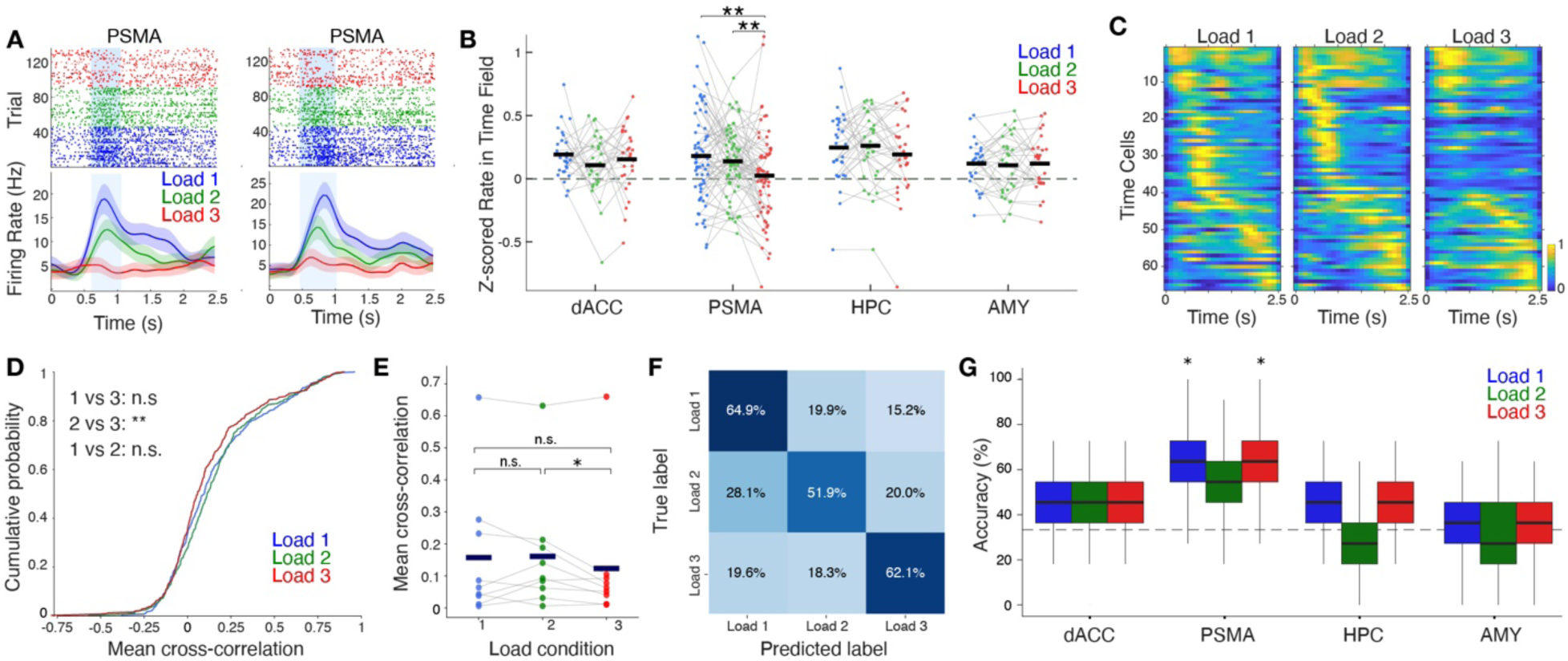
| Memory load modulates temporal coding by PSMA time-cells. **A.** Raster plots and mean± SEM firing rates of example time cells across the 3 Load conditions. t=0 is onset of the maintenance period. Blue bars mark time fields as before. **B.** Mean z-scored within-field firing rates of each time cell, averaged across trials of each load. Two-sided paired-sample t-tests. dACC: Load 1 vs 2, p = 0.2187; Load 1 vs 3, p = 0.5360; Load 2 vs 3, p = 0.4826. PSMA: Load 1 vs 2, p = 0.3106; Load 1 vs 3, p = 0.0089; Load 2 vs 3, p = 0.0078. HPC: Load 1 vs 2, p = 0.8035; Load 1 vs 3, p = 0.3771; Load 2 vs 3, p = 0.0887. AMY: Load 1 vs 2, p = 0.7486; Load 1 vs 3, p = 0.9935; Load 2 vs 3, p = 0.7966. **C.** Average firing rates of PSMA time-cells across trials of each load, ordered by time field over all trials. **D.** Cumulative distributions of pairwise time cell cross-correlations per trial for each load. (Kolmogorov–Smirnov tests: Load 1vs2, p = 0.276; Load 1vs3, p = 0.128; Load 2vs3, p = 0.00278). **E.** Per-patient mean cross-correlations across memory loads (one-sided paired-sample t-test: Load 1 > 2 p = 0.582.; Load 2 > 3, p = 0.031; Load 1 > 3, p = 0.128). **F.** Confusion matrix of accuracy in decoding memory load from PSMA time cells using a three-way linear support-vector machine (SVM) for each true load (rows) versus predicted load. **G**. Load-decoding with 3-way linear SVM models trained and tested on time cells of each recorded brain region. Empirical t-test (actual accuracies from 1000 monte-carlo splits of train/test trials against same number of null decoding accuracies). Per corresponding load: dACC: p = 0.231, 0.266, 0.129; PSMA: p = 0.03, 0.112, 0.049. HPC: p = 0.393, 0.455, 0.261. AMY: p = 0.361, p = 0.472, p = 0.273.

To compare PSMA time cell coordination, we computed average pairwise cross-correlations, as before, separately for trials of each load condition. Cumulative cross-correlations were significantly lower in Load 3 than Load 2 trials, both over pooled trials (Fig. 3D) and at per-patient level (Fig. 3E), whereas Load 1 was not significantly different to the other conditions.

To further quantify load-modulation effects on a population level, we trained linear SVM classifiers to decode memory load based on time-cell firing during the maintenance period. Only the decoder trained and tested on PSMA had overall above chance performance (empirically p = 0.008) in classifying Load 1 and Load 3 trials but not Load 2 (Fig. 3G), confirming distinct PSMA time cells activations between load conditions (Fig. 3F-G). Importantly, we repeated out analyses using only correct trials within each Load, to exclude any potential effects of successful WM, but our findings remained unaffected; namely reduced within-field modulation in Load 3 trials (two-sided t-test, Load 1 vs 2 p=0.195, Load 2 vs 3 p = 0.0127, Load 1 vs 3 p =0.00524), diminished cross-correlations (Kolmogorov–Smirnov tests: Load 1 vs 2, p = 0.255; Load 1 vs 3, p =0.167; Load 2 vs 3, p = 0.0114), and above-chance load decoding (p = 0.008). This confirms that the apparent weakening of temporal modulation under Load 3 cannot be attributed to the higher error rate observed for Load 3 trials (which are more difficult).

In conclusion, like the persistent spiking of PSMA maintenance cells, the temporal modulation of PSMA time cells during WM maintenance reflects the number of items carried in memory.

### Cue-Selective Time Cells During WM Maintenance

So far, time cells have been detected based on maintenance activity across all trials. However, time cells in rodents, firing during a post-cue delay period, can be specific to the preceding cue, thereby encoding WM content^17^. We thus examined whether time-cell activation during the maintenance period could be selective for the identity of the images currently held in WM.

To associate maintenance activation with specific preceding images, rather than combinations of images, we focused on Load 1 trials only. We found ‘cue-selective’ time cells (n = 91 total) with significant temporally tuned activation during maintenance, only after a specific (‘preferred’) image was presented, and no significant tuned activation after all other four images used in the task (Fig. 4A; See Methods). Sorting cue-selective time cells by their field timepoint yielded a activation sequence in their corresponding preferred trials (maintenance after the preferred image in Load 1 trials), tiling the maintenance period, that was absent in non-preferred trials (maintenance after non-preferred image; Fig. 4B). This sequence was validated across independent halves of preferred trials demonstrating its stability across trials (Fig. 4C). Image-selective time cells were more common than expected by chance in all five recorded brain areas (Fig. 4D; dACC: 13/171, 7.6%; PSMA: 32/250, 12.8%; hippocampus: 14/190, 7.4%; amygdala: 26/259, 10.0%; vmPFC: 6/32, 18.8%). Across patients, the mean proportion of cue-selective time cells was 11.25% ± 1.75% (mean ± SEM).

**Fig. 4.**
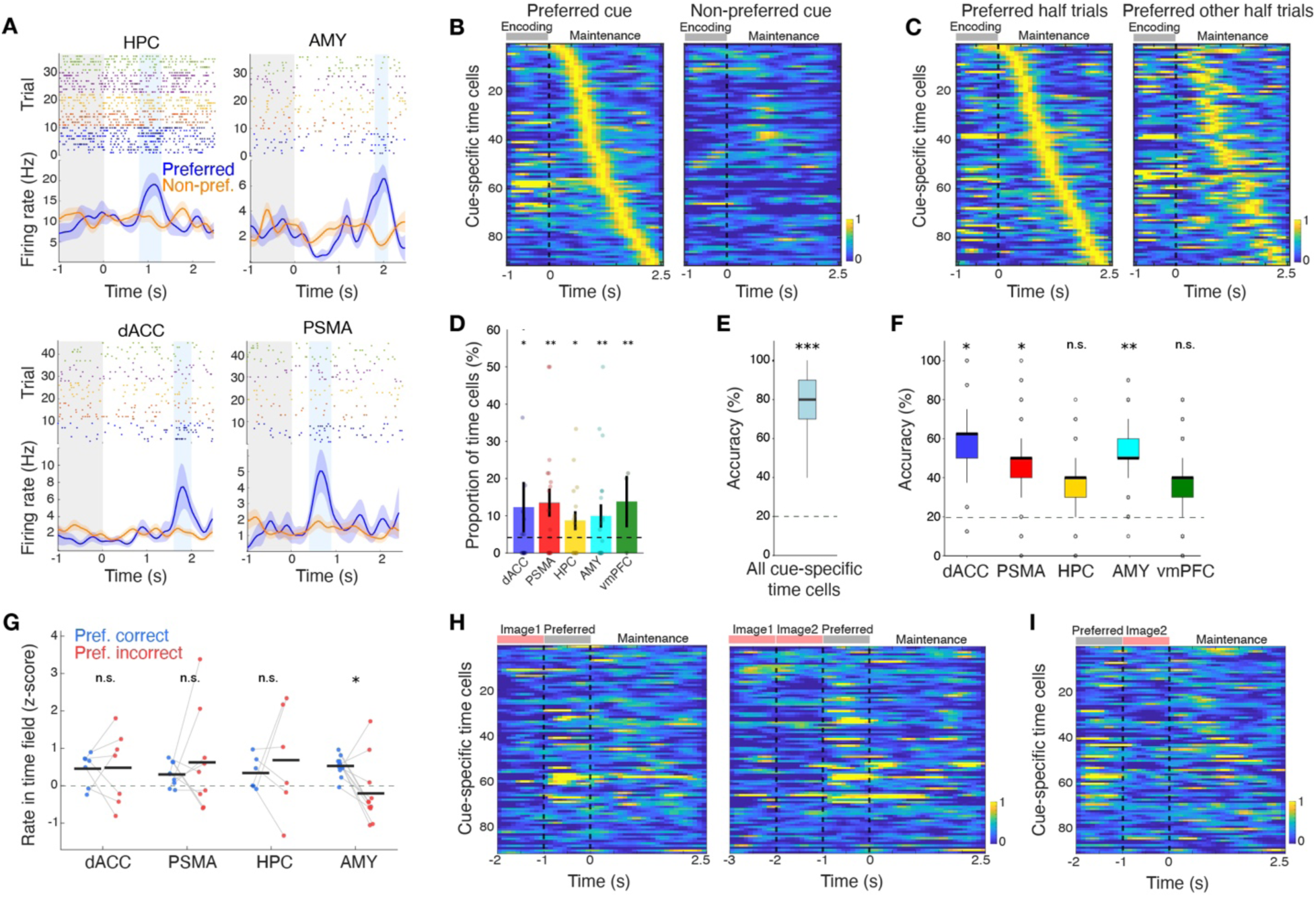
| Cue-selective time-cells in Load 1 trials encode cue identity during WM. **A.** Raster plots and mean± s.e.m z-scored firing rates across the encoding (gray bar) and maintenance phase, from example cue-selective time cells in Load-1 trials where the neuron’s preferred cue (blue) or non-preferred cues were shown (orange). Colored dots in the raster plot indicate spikes during and after each one of the 5 images, with blue always indicating the preferred one. **B.** Normalized average firing rates of pooled cue-selective time cells across encoding and maintenance phase in their corresponding preferred (left) and non-preferred trials (right). Neurons are ordered by their time fields. Dashed line: Maintenance onset. **C.** Normalized average firing rates of pooled cue-selective time cells across randomly selected 50% of preferred trials (left) and the other 50% (right). In both plots, neurons are sorted by firing rate peak times in the first set of trials. **D.** Mean± s.e.m. percentage of cue-selective time cells per brain region in each patient (dots). Asterisks indicate significance of regional cell counts against an empirical null distribution (See methods; empirical p value, p_dACC_ = 0.0399, p_PSMA_ = 0.002, p_HPC_ = 0.022, p_AMY_ = 0.002, p_vmPFC_ = 0.004). One outlier data point in dACC was omitted for plotting clarity. **E.** Accuracy of decoding cue identity in Load1 trials based on maintenance-phase within-field z-scored firing of cue-selective time cells, with a 5-way linear SVM. Dashed line: chance baseline. Empirical p = 0 against null distribution. **F.** Mean accuracy of 5-way SVM cue decoding in Load 1 trials, using only cue-selective time cells from each region (1000 splits of training/testing trials). Empirical p-values against chance baseline respectively: p_dACC_ = 0.0315, p_PSMA_ = 0.0342, p_HPC_ =0.1254, p_AMY_ = 0.008, p_vmPFC_ = 0.119.(See Methods). **G.** Within-field z-scored firing rates of cue-selective time cells in preferred correct versus preferred incorrect Load 1 trials (n = 36 neurons in total from patients with at least one incorrect trial; two-sided paired-sample t-test). dACC (n = 7 cells, p = 0.529), pre-SMA (n = 9, p = 0.739), amygdala (n = 11, p = 0.012), and hippocampus (n = 6, p = 0.692). **H.** Same as **B,** with same cells ordering, but for Load 2 trials where preferred image was preceded by one (left) or two (right) non-preferred image(s). **I.** Same for trials where preferred image was followed by a non-preferred image.

To assess whether cue-selective time cells reliably encoded memory content in single trials, we decoded the identity of the image held in WM in Load 1 trials from the activation of pooled cue-selective time cells using a 5-way linear SVM classifier (see Methods). Average decoding accuracy was significantly higher than chance of 20% (Fig. 4E), confirming that time cells as a group retain the memory of the preceding image throughout maintenance. When pooling cells separately per brain region, decoding accuracy was significantly higher than chance in dACC, PSMA and amygdala (Fig. 4F). Moreover, correct Load 1 trials yielded significantly higher spike tuning of cue-selective time cells in the Amygdala compared to incorrect trials, but not in the other brain regions (Fig. 4G).

Finally, we tested whether the selectivity of cue-selective time cells was linked to their corresponding preferred image, as a visual cue, or a specific memorandum. We reasoned that if these time cells were triggered after their preferred image as a visual cue, then they would also be triggered after the same image in Load 2 trials, yielding similar sequences when the preferred image was shown last, or time-shifted sequences when the preferred image was shown first. However, cue-selective time cells did not exhibit temporal tuning in higher load trials, when their preferred image was preceded by 1 or 2 non-preferred images (Fig 4H) or when the preferred image was followed by a non-preferred image (Fig. 4I). Similarly, we could detect time cells that fired after a particular final image in Load 2 trials (n = 64), or in Load 3 trials (n = 49) but the emerging sequences did not hold in other Load conditions when the same image was last.

Together, these findings revealed cue-selective time cells that are not passively triggered by an image itself but by a specific memorandum. Their activation encodes this content more efficiently in frontal regions and in the amygdala.

### Complementary Persistent Firing-Based and Sequence-Based Codes during WM Maintenance

Previous studies have focused on visually selective ‘concept cells’, whose firing rate during a visual stimulus increases only for their preferred image. Concept cells continue to fire persistently across the maintenance period if the preferred image is held in memory, thereby retaining information in WM^6,33,34^. We thus next asked whether there was a relationship between concept cells, time cells, and cue-specific time cells.

We identified concept cells with selective activation during the presentation of a given image across loads, when the image was shown first (see Methods; n = 119). Concept cells were mostly observed in Amygdala (71/259 units), followed by hippocampus (23/190), PSMA (12/250), dACC (8/171) and vmPFC (5/32). As expected, concept cells persistently spiked during the ensuing maintenance period when their preferred stimulus was held in WM in addition to being selective during presentation of the stimulus itself (Fig. 5A, blue). In contrast, cue-specific time cells respond selectively only during the maintenance period, but not during the encoding period (Fig. 5A, orange). The two cell populations were largely distinct: only 21 of 119 concept cells (17.6%) were also cue-specific time cells (Fig. 5B)

**Fig. 5.**
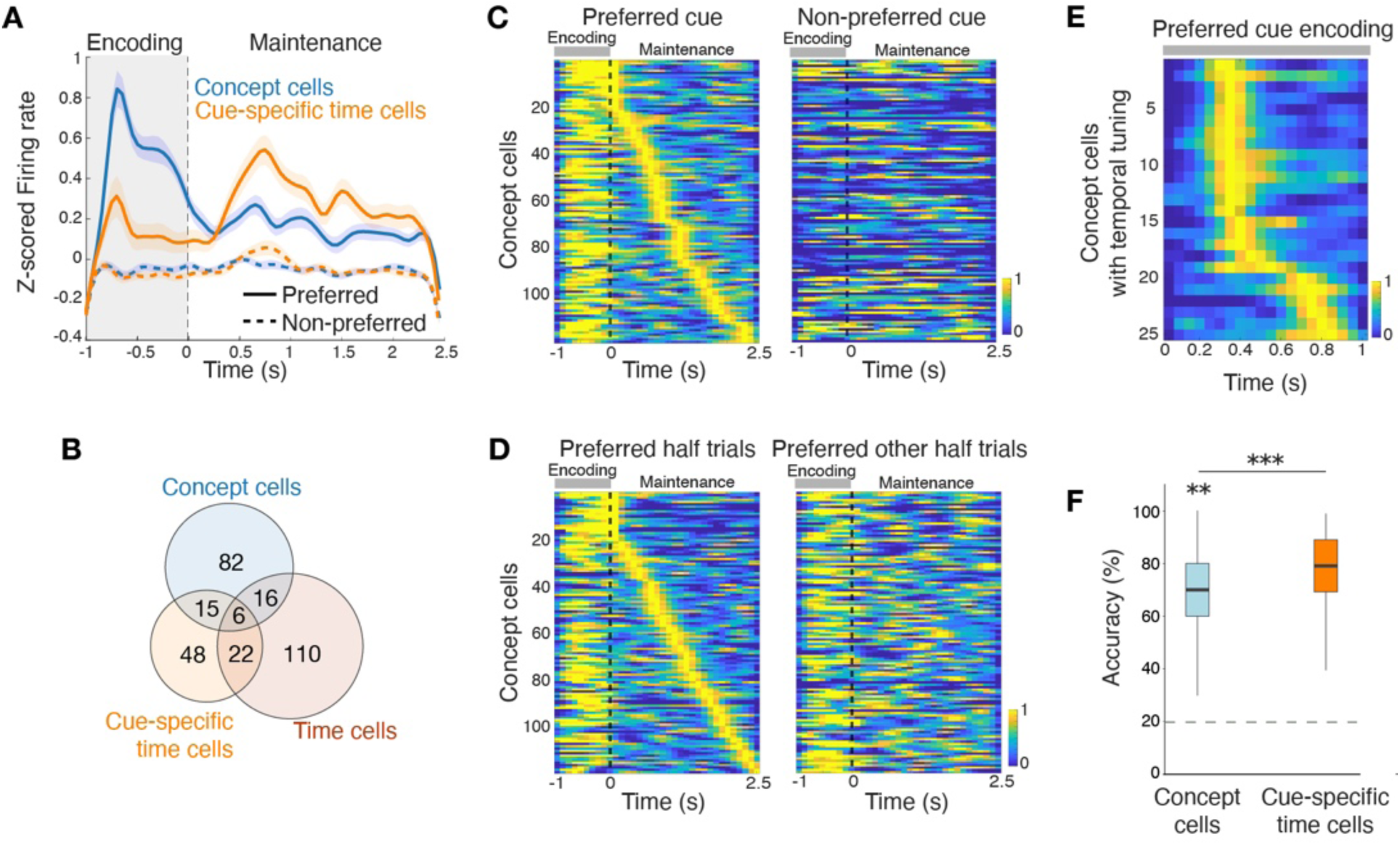
| Parallel retention of cue information by cue-selective time cells and persistently firing concept cells. **A.** Mean z-scored firing rates of concept cells and cue-selective time cells, throughout encoding and maintenance, during Load-1 trials with preferred vs non-preferred images encoded. **B.** Venn diagram showing the overlapping between different types of identified neurons in this study. **C.** Average normalized firing rates of concept cells (n = 119) across Load 1 preferred-image trials (left) and non-preferred trials (right), ordered by peak latency during maintenance in preferred trials. **D.** Same for random half of preferred trials (left) versus the other half (right), with neurons sorted based on the first group’s maintenance firing peaks. **E.** Average firing rates of concept cells with temporally tuned response to preferred images (n = 25 cells) during the preferred image presentation in Load 1 trials. **F.** Blue: Accuracy of decoding cue identity in Load1 trials based on average firing rates of concept cells across maintenance, with a 5-way linear SVM. Orange: Same for SVM accuracy from cue-selective time cells (as in Fig 4F). Dashed line: chance baseline. Empirical p-value relative to label-shuffled null for concept cell-based SVM = 0.003. Cue-selective time cells yielded significantly higher decoding accuracy; Wilcoxon test p = 1.37 * 10^-36^.

Persistent spiking of concept cells during maintenance was specific to their preferred image (Fig. 5C). Even though transient spiking peaks could be sorted to form a sequence (by definition), they lacked consistent temporal tuning, demonstrated by comparing two sets of Load 1 trials (Fig. 5D). Concept cells also did not exhibit temporal tuning during encoding: Only 25/119 (21%) of concept cells exhibited a time field during encoding, when their preferred image was shown in Load 1 trials (See Methods; p < 0.05). In addition, the fields of these temporally modulated concept cells were concentrated within 500 ms from image onset, rather than forming a sequence, likely encoding image onset rather than image presentation time (Fig. 5E). Image identity could be decoded from the average activity of concept cells during maintenance, using a five-way SVM classifier, confirming memory encoding through their persistent spiking. However, decoding accuracy by concept cells was significantly lower than by cue-selective time cells (Fig. 5F). This difference remained significant after excluding neurons that were shared between the two populations (Wilcoxon test, p = 2.5x10^-105^).

Collectively, these results show that concept cells and cue-specific time cells provide parallel codes for WM encoding, but time cells provide more efficient information encoding in WM.

## Discussion

Our results demonstrate that the human brain internally generates time-cell sequences during WM maintenance. These sequences tile the delay interval, emerge in the absence of explicit timing demands, and predict both the accuracy and quality of memory performance. Cue-specific time cells carry explicit mnemonic information, providing a temporal code for WM maintenance, complementary to sustained spiking by concept cells. Together, our findings reveal for the first time an interplay of persistent and sequential dynamics supporting human WM.

### Time-cell sequences as a human WM mechanism

Theoretical^35,36^, experimental^37^, and computational^26,38^ work has illustrated the importance of self-generated sequential activity in cognition. Time-cell sequences have been extensively characterized in rodents (in hippocampus, striatum, parietal, entorhinal and prefrontal cortex) where they provide a temporal scaffold for memory across delays and discontiguous events^8,9,22,39–41^. Here, we show that analogous sequences exist in the human brain during a non-spatial WM task. Unlike previous human studies where temporal modulation reflected external structure or explicit temporal demands^27,28^, the Sternberg task requires only memorandum identity maintenance. The emergence of time-cell sequences under these conditions demonstrates that temporal coding can be internally generated in humans to support WM.

Notably, similar sequences were absent during continuous video viewing^30^, where neural activity instead tracked event boundaries. This dissociation suggests that the brain flexibly recruits internal versus externally anchored temporal representations depending on task structure. When external stimuli provide a temporal framework to-be-remembered (e.g. word lists^28^), temporal coding may be driven by boundaries or similar specific temporal slots. When information must be actively maintained over short delays, as in the task used here or in a delayed non-match-to-sample task used in mice^17,18^, internally generated sequences become dominant.

Our results therefore provide experimental support for temporal context models ^20,23^, positing that a drifting temporal code enables the brain to bind ‘what’ to ‘when’. They also support the postulation that time-cell sequences provide a domain-general “sequence engine” for organizing cognition^19^. In WM tasks such as the present paradigm, maintaining the temporal progression of the delay is essential for sustaining internal task state^42,43^. Thus, time-cell dynamics provide the temporal scaffold that supports maintenance, complementing models that emphasize persistent rate coding^6^, and theta–gamma phase–amplitude coupling^44^.

### Functional specialization across brain regions

Our data reveal region-specific contributions of time-cell sequences to WM. In the hippocampus and dorsal anterior cingulate cortex, stronger and more coordinated time-cell activity predicted successful memory maintenance. One interpretation is that precise and globally aligned sequences provide a temporal basis to tether their stimulus-specific codes and to maintain internal task state. When this basis becomes noisy, the fidelity of time cell sequence degrades, and behaviour suffers.

In contrast, time-cell activity in pre-supplementary motor area was selectively modulated by memory load and predicted response speed (a measure of memory quality in Sternberg-type tasks^31,32^) rather than accuracy. This mirrors prior findings that persistent PSMA spiking scales with WM load^45^ and suggests that temporal coding in PSMA reflects the computational demands imposed by multiple memoranda.

Notably, single neurons in the amygdala have been known to encode high-level visual and affective stimuli^46,47^, and to predict successful associative memory and recognition performance^48,49^. Similarly, persistent activity in the amygdala is often seen during WM tasks with visual stimuli as memoranda^6,50^. Consistent with this, amygdala time cells in our task showed strong cue-encoding and reduced tuning in incorrect trials.

Together, these results indicate that time-cell sequences are differentially tuned across brain regions to support distinct aspects of WM.

### Cue-selective time cells and complementary mnemonic codes

A key finding of this study is the identification of cue-selective time cells in the human brain. These neurons fired at specific moments during the maintenance period only when a particular item was held in memory, encoding both item identity and elapsed time. Importantly, these responses were not simply triggered by visual input but depended on the mnemonic status of the item, consistent with rodent studies showing learning-dependent emergence of odor-specific time cells^17^.

Cue-selective time cells formed a coding scheme distinct from classical concept cells. Whereas concept cells encoded item identity through persistent firing, cue-selective time cells provided a temporally structured, distributed code. The two populations overlapped minimally and supported memory decoding through different mechanisms, suggesting parallel and complementary mnemonic representations that may be synergistic, providing the brain with multiple readout strategies for memory content. This dual-code framework aligns with recent theoretical work proposing that neural systems simultaneously encode content and temporal structure via multiplexed or orthogonal dynamics^26^.

### Future directions

Looking ahead, future work will be needed to determine whether human time cells can code temporal information across longer timescales of minutes^51^ to days^52^, or how they interact with oscillatory dynamics and boundary representations in naturalistic settings; questions that will be crucial for linking WM mechanisms to episodic memory across contexts. Also, future studies using repeated stimuli combinations in Load 2 and 3 trials will establish if cue-specific time cells can indeed encode more complex specific memoranda. Understanding how these temporal codes break down in neurological and psychiatric disorders may also provide insight into the mechanisms underlying memory dysfunction.

## Methods

### Datasets Description

#### Sternberg WM Task

This dataset includes human single-neuron activity recorded during an initial screening task and a subsequent modified Sternberg WM task. Data were collected from 21 patients (aged 17–71) undergoing intracranial monitoring for drug-resistant epilepsy, using hybrid macro-micro depth electrodes targeting the medial temporal lobe (hippocampus, amygdala) and medial frontal regions (anterior cingulate cortex, pre-supplementary motor area, ventromedial prefrontal cortex). Across 41 sessions, 1,809 neurons were isolated in total: 907 during screening and 902 during the task. Continuous broadband recordings (0.1–9,000 Hz) were sampled at 32 kHz, with spike detection and sorting performed using a semi-automated template-matching algorithm and rigorous isolation quality metrics.

In the screening task, participants viewed 54–64 unique images (e.g., people, animals, scenes) presented six times each in randomized order for 1 s with variable interstimulus intervals. Periodic catch questions (e.g., “Was the last image an animal?”) ensured attention. Single-neuron selectivity was assessed online to identify concept cells showing stimulus-specific firing. The five images eliciting the strongest selective responses were then used as memoranda in the task. During each trial, after a variable fixation period lasting between 0.9 and 1 seconds, 1–3 images were shown sequentially (each lasting for 1 second), followed by a 2.5–2.8 s maintenance period and a probe requiring a match/non-match decision.

For details please refer to relevant past studies^6,29^.

#### Video-clip Task

Participants viewed a series of 90 silent video clips and were instructed to encode each clip as thoroughly as possible. Each trial commenced with a variable baseline fixation period lasting between 0.9 and 1.1 seconds (sampled from a uniform distribution), during which a central fixation cross reminded participants to maintain gaze at the center of the screen. This was followed by the presentation of a video clip containing one of three boundary types: (1) no boundaries (NBs), consisting of a continuous movie shot with a virtual midpoint boundary used for analytic purposes; (2) soft boundaries (SBs), representing within-movie scene changes, with one to three SBs randomly interspersed throughout the clip; or (3) hard boundaries (HBs), indicating a transition to a scene from a different movie, consistently positioned 4 seconds after clip onset. Illustrative examples of SBs and HBs are provided in Fig.S1A. To ensure engagement, a yes/no question related to clip content (e.g., “Is anyone in the clip wearing glasses?”) was randomly interleaved every four to eight trials.

For subject information, spike sorting and metrics, please refer to the studies^30^.

#### Selection of Neurons

To identify time cells, we employed a permutation-based approach. Each unit’s observed firing rate vector, averaged across trials after binning (bin width = 100ms), Gaussian smoothing (σ = 150ms), and Z-scoring, was compared against a null distribution generated by randomly circularly shifting spike times within each trial. For each permutation, spike trains were reprocessed identically (binned, smoothed, Z-scored, and averaged) to yield a permuted firing rate vector. The empirical p-value was computed by comparing the maximum value of the observed vector to the distribution of maxima from the permuted vectors. Units showing significant temporal modulation (p < 0.05) were classified as time cells. The time field of each time cell was defined as the bin with the maximum Z-scored firing rate.

To identify cue-selective time cells, maintenance-phase spike counts were smoothed by the same Gaussian kernel on a per-trial basis (only Load 1 trials), and the resulting traces were averaged across trials of each image, to obtain an image-specific firing-rate profile. For each image, we quantified the peak amplitude of this profile. Only images that met the minimum trial-count criterion (7 trials) were considered as candidate images for that unit. To control for multiple comparisons across image categories within a unit, we used a permutation-based max-T procedure. For each of 1,000 permutations, spike trains were randomly circularly shifted within each trial and image labels were randomly reassigned across trials while preserving the number of trials per image. For each permuted dataset, we recomputed the image-specific firing-rate profiles as for the observed data and, for each image, extracted the peak of the mean Z-scored firing-rate over time. We then took the maximum of these peak values across images, yielding one unit-level maximum peak firing rate for that permutation. The same unit-level statistic (maximum image-wise peak firing rate) was computed from the observed data. An empirical p-value was obtained as the proportion of permutations in which the permuted unit-level maximum was greater than or equal to the observed maximum. Units with p < 0.05 were classified as cue-selective time cells. Final-image-selective time cells in Load 2 and 3 conditions were identified using the same algorithm, with trials segregated by the last-image identity.

Single units were classified as concept cells if their firing rates significantly varied with image identity during the encoding period. We computed a one-way ANOVA (F-statistic) on mean firing rates in a 200–1,000 ms window post-stimulus onset (α = 0.05). Cells passing this test were further required to show a significantly higher response to the preferred image than all others (permutation t-test, p < 0.05). As previously demonstrated, this procedure ensured robust selectivity for visual stimuli used in subsequent WM trials^6^.

To identify the subpopulation of concept cells with temporally modulated response to preferred images in load 1 trials, we ran the same permutation t-test for identifying time cells, but during the first second of the preferred image presentation. Concept cells with significant temporal tuning were identified (p < 0.05).

#### Chance level of Cell Selection

To estimate a false discovery rate for both time cells and cue-specific time cells, we performed permutation tests. For each neuron, spike times were circularly shifted randomly, after which we reapplied the previously described detection procedure to identify time cells and compute their proportion within each region. This randomization was repeated 1,000 times to generate a null distribution of time cell counts.

An analogous procedure was used to evaluate cue-specific time cells. In this case, after shuffling spike timing, the image labels associated with load-1 trials were also randomly reassigned, yielding a permuted dataset and a corresponding null distribution for cue-specific time cell counts. Statistical significance was determined by comparing the observed number of time cells or cue-specific time cells in each region in the real data to their respective null distributions of cell yields derived from the shuffled trials.

#### Cross-Correlation Analysis of Time Cell Activity

To assess the temporal coordination of time cell firing, we computed cross-correlations between all time cell pairs during individual trials. Prior to correlation analysis, we circularly shifted each neuron’s firing rate vector so that its time field was centered within the trial window. For each trial, we computed the Pearson correlation coefficient between all neuron pairs and averaged these values to obtain a trial-level mean cross-correlation.

To quantitatively evaluate how last-image-specific time cells defined in one load(reference load) preserve the sequence in other load conditions, for each of the other loads, we computed Pearson correlation coefficient between the average firing rate of trials of the load versus trials of the reference load during the maintenance session following an encoding series ending with its preferred image identified.

#### Resampled Cross-Correlation Distribution Analysis

To evaluate whether correct trials exhibited stronger time cells synchrony than incorrect trials, we compared cross-correlation distributions using a resampling approach, considering the difference between the sample size of correct versus error trials. We first computed the mean cross-correlation across all incorrect trials. Next, we repeatedly sampled an equal number of correct trials (without replacement) and computed the mean cross-correlation for each resampled set. This process was iterated 1,000 times to generate a distribution of mean cross-correlations for correct trials. A significance test was performed by empirically calculating p-value as the fraction of resampled correct trial means falling below the incorrect trial mean.

#### Population Decoding

##### Support Vector Machine

To assess whether ensemble neuronal activity carried decodable information about cognitive variables of interest (e.g., WM load, reaction time, or stimulus identity), we implemented a pseudopopulation-based decoding framework using linear Support Vector Machines (SVMs). This approach constructs a virtual population by aggregating neurons recorded across multiple sessions and subjects, enabling robust population-level inference even when simultaneous recordings are not available.

Per time cell, trial-averaged z-scored or raw firing rates were extracted over their time field. To equalize trial counts across conditions, firing rates were downsampled to the minimum number of trials observed per condition across the dataset. For each pairwise decoding comparison, we constructed labeled pseudopopulation matrices by stacking firing rate vectors across neurons. These matrices were then randomly partitioned into training and testing sets in a repeated cross-validation loop (1,000 iterations), with decoder performance quantified as classification accuracy on held-out trials.

To establish statistical significance, a nonparametric permutation test was conducted in each iteration by randomly shuffling condition labels in the training data. This generated a null distribution of decoding accuracy, against which the actual decoding performance was compared to derive empirical p-values.

##### Naive Bayes Decoding

To assess the capacity of neuronal populations to encode temporal information during WM maintenance, we applied population decoding using a Gaussian Naive Bayes classifier. Neural activity was binned into 100ms windows over the 2.5 s delay period and organized into matrices (time bins × trials × neurons). To ensure balanced sampling, the number of trials was truncated to match the neuron with the fewest recorded trials.

The data were split into training and test sets by partitioning trials (70% training, 30% testing), maintaining the temporal structure of each trial. A separate decoder was trained for each population of interest—time cells and other cells—using the binned firing rates as input features and the corresponding time bin as the target label. Decoding performance was quantified by computing the absolute difference between the true and predicted time bins, scaled to seconds.

To establish a baseline for chance-level decoding, we performed a permutation test in which the temporal labels of the test set were randomly shuffled 1,000 times. The classifier’s predictions on the shuffled labels provided a distribution of baseline decoding errors for comparison.

Mean decoding errors and standard errors were computed for each time bin and compared across conditions. Pairwise statistical comparisons between time cells, non–time cells, and baseline conditions were performed using two-tailed Welch’s t-tests. All analyses were implemented in Python using scikit-learn and our custom scripts for data preprocessing, model fitting, and statistical testing.

#### Per-Patient Naive Bayes Decoding

To examine how population coding of temporal information varies with behavioral performance, we conducted a per-patient decoding analysis comparing correct and incorrect trials. For each patient, spiking activity was binned across the delay interval and organized into matrices with dimensions (time bins × trials × neurons). Neuronal ensembles were included if they have five or more neurons to ensure sufficient population size.

Trials were categorized as correct or incorrect based on trial outcome labels. To control for trial count imbalance, the test set for correct trials was matched to the number of incorrect trials for each patient. A Gaussian Naive Bayes classifier was trained using only correct trials and tested on both incorrect trials and held-out correct trials of the same size. Decoding performance was quantified as the absolute difference between the predicted and true time bin labels, multiplied by the bin width to yield decoding errors in seconds. In Fig. S3, for each region, decoding was performed once using all neurons and once with neurons from that region removed, using identical train–test splits, on patients with ≥5 total time cells, ≥1 time cell in the target region, ≥1 time cell remaining after dropout, and at least 1 incorrect trial.

We also conducted per-patient decoding analysis to compare time cells versus other cells (Fig. 1J), with train test split ratio equal to 0.3.

## Acknowledgments

We would like to thank Drs. Kathrin Duncan and Kaori Takehara for valuable feedback on the analysis. This work was supported by the New Frontiers in Research Fund (NFRFE-2024-00388) and the BRAIN initiative (U01NS103792, U01NS117839).

## References

1. Baddeley, A. Working Memory. Science 255, 556–559 (1992).

2. D’Esposito, M. & Postle, B. R. The Cognitive Neuroscience of Working Memory. Annual Review of Psychology 66, 115–142 (2015).

3. Goldman-Rakic, P. S. Cellular basis of working memory. Neuron 14, 477–485 (1995).

4. Fuster, J. M. & Alexander, G. E. Neuron activity related to short-term memory. Science 173, 652–654 (1971).

5. Constantinidis, C. et al. Persistent Spiking Activity Underlies Working Memory. J. Neurosci. 38, 7020–7028 (2018).

6. Kamiński, J. et al. Persistently active neurons in human medial frontal and medial temporal lobe support working memory. Nat Neurosci 20, 590–601 (2017).

7. Pastalkova, E., Itskov, V., Amarasingham, A. & Buzsáki, G. Internally Generated Cell Assembly Sequences in the Rat Hippocampus. Science 321, 1322–1327 (2008).

8. MacDonald, C. J., Lepage, K. Q., Eden, U. T. & Eichenbaum, H. Hippocampal “Time Cells” Bridge the Gap in Memory for Discontiguous Events. Neuron 71, 737–749 (2011).

9. Harvey, C. D., Coen, P. & Tank, D. W. Choice-specific sequences in parietal cortex during a virtual-navigation decision task. Nature 484, 62–68 (2012).

10. Schuck, N. W. & Niv, Y. Sequential replay of nonspatial task states in the human hippocampus. Science 364, eaaw5181 (2019).

11. Miller, E. K. & Cohen, J. D. An Integrative Theory of Prefrontal Cortex Function. Annual Review of Neuroscience 24, 167–202 (2001).

12. Romo, R., Brody, C. D., Hernández, A. & Lemus, L. Neuronal correlates of parametric working memory in the prefrontal cortex. Nature 399, 470–473 (1999).

13. Pesaran, B., Pezaris, J. S., Sahani, M., Mitra, P. P. & Andersen, R. A. Temporal structure in neuronal activity during working memory in macaque parietal cortex. Nat Neurosci 5, 805–811 (2002).

14. Machens, C. K., Romo, R. & Brody, C. D. Functional, But Not Anatomical, Separation of “What” and “When” in Prefrontal Cortex. J. Neurosci. 30, 350–360 (2010).

15. Murray, J. D. et al. Stable population coding for working memory coexists with heterogeneous neural dynamics in prefrontal cortex. Proc Natl Acad Sci U S A 114, 394–399 (2017).

16. Brody, C. D., Hernández, A., Zainos, A. & Romo, R. Timing and Neural Encoding of Somatosensory Parametric Working Memory in Macaque Prefrontal Cortex. Cereb Cortex 13, 1196–1207 (2003).

17. Taxidis, J. et al. Differential Emergence and Stability of Sensory and Temporal Representations in Context-Specific Hippocampal Sequences. Neuron 108, 984–998.e9 (2020).

18. Dorian, C. C., Taxidis, J., Buonomano, D. V. & Golshani, P. Hippocampal sequences represent working memory and implicit timing. Cell Reports 44, (2025).

19. Buzsáki, G. & Tingley, D. Space and Time: The Hippocampus as a Sequence Generator. Trends in Cognitive Sciences 22, 853–869 (2018).

20. Eichenbaum, H. Time cells in the hippocampus: a new dimension for mapping memories. Nat Rev Neurosci 15, 732–744 (2014).

21. Ma, M. et al. Sequential activity of CA1 hippocampal cells constitutes a temporal memory map for associative learning in mice. Current Biology 34, 841–854.e4 (2024).

22. Kraus, B. J., Robinson, R. J., White, J. A., Eichenbaum, H. & Hasselmo, M. E. Hippocampal “Time Cells”: Time versus Path Integration. Neuron 78, 1090–1101 (2013).

23. Ranganath, C. & Hsieh, L.-T. The hippocampus: a special place for time. Annals of the New York Academy of Sciences 1369, 93–110 (2016).

24. Howard, M. W. et al. A Unified Mathematical Framework for Coding Time, Space, and Sequences in the Hippocampal Region. J. Neurosci. 34, 4692–4707 (2014).

25. Rolls, E. The mechanisms for pattern completion and pattern separation in the hippocampus. Front. Syst. Neurosci. 7, (2013).

26. Zhou, S., Seay, M., Taxidis, J., Golshani, P. & Buonomano, D. V. Multiplexing working memory and time in the trajectories of neural networks. Nat Hum Behav 7, 1170–1184 (2023).

27. Schonhaut, D. R., Aghajan, Z. M., Kahana, M. J. & Fried, I. A neural code for time and space in the human brain. Cell Reports 42, 113238 (2023).

28. Umbach, G. et al. Time cells in the human hippocampus and entorhinal cortex support episodic memory. Proceedings of the National Academy of Sciences 117, 28463–28474 (2020).

29. Kyzar, M. et al. Dataset of human-single neuron activity during a Sternberg working memory task. Sci Data 11, 89 (2024).

30. Zheng, J. et al. Neurons detect cognitive boundaries to structure episodic memories in humans. Nat Neurosci 25, 358–368 (2022).

31. Sternberg, S. High-speed scanning in human memory. Science 153, 652–654 (1966).

32. Ratcliff, R. A theory of memory retrieval. Psychological Review 85, 59–108 (1978).

33. Daume, J. et al. Persistent activity during working memory maintenance predicts long-term memory formation in the human hippocampus. Neuron 0, (2024).

34. Kornblith, S., Quian Quiroga, R., Koch, C., Fried, I. & Mormann, F. Persistent Single-Neuron Activity during Working Memory in the Human Medial Temporal Lobe. Current Biology 27, 1026–1032 (2017).

35. Hayon, G., Abeles, M. & Lehmann, D. A Model for Representing the Dynamics of a System of Synfire Chains. J Comput Neurosci 18, 41–53 (2005).

36. Buzsáki, G. Two-stage model of memory trace formation: A role for “noisy” brain states. Neuroscience 31, 551–570 (1989).

37. O’Keefe, J. & Dostrovsky, J. The hippocampus as a spatial map. Preliminary evidence from unit activity in the freely-moving rat. Brain Research 34, 171–175 (1971).

38. Lin, D., Huang, A. Z. & Richards, B. A. Temporal encoding in deep reinforcement learning agents. Sci Rep 13, 22335 (2023).

39. Akhlaghpour, H. et al. Dissociated sequential activity and stimulus encoding in the dorsomedial striatum during spatial working memory. eLife 5, e19507 (2016).

40. Tiganj, Z., Jung, M. W., Kim, J. & Howard, M. W. Sequential Firing Codes for Time in Rodent Medial Prefrontal Cortex. Cereb Cortex 27, 5663–5671 (2017).

41. Takehara-Nishiuchi, K. Neuronal ensemble dynamics in associative learning. Current Opinion in Neurobiology 73, 102530 (2022).

42. Howard, M. W., Shankar, K. H., Aue, W. R. & Criss, A. H. A distributed representation of internal time. Psychological Review 122, 24–53 (2015).

43. Buzsáki, G. Time, space, memory and brain–body rhythms. Nature Reviews Neuroscience 1–18 (2025).

44. Daume, J. et al. Control of working memory by phase–amplitude coupling of human hippocampal neurons. Nature 629, 393–401 (2024).

45. Owen, A. M. et al. Functional organization of spatial and nonspatial working memory processing within the human lateral frontal cortex. Proceedings of the National Academy of Sciences 95, 7721–7726 (1998).

46. Mormann, F. et al. A category-specific response to animals in the right human amygdala. Nat Neurosci 14, 1247–1249 (2011).

47. Kreiman, G., Koch, C. & Fried, I. Imagery neurons in the human brain. Nature 408, 357–361 (2000).

48. Rutishauser, U., Schuman, E. M. & Mamelak, A. N. Activity of human hippocampal and amygdala neurons during retrieval of declarative memories. Proceedings of the National Academy of Sciences 105, 329–334 (2008).

49. Rutishauser, U. et al. Representation of retrieval confidence by single neurons in the human medial temporal lobe. Nature Neuroscience 18, 1041–1050 (2015).

50. Kamiński, J., Brzezicka, A., Mamelak, A. N. & Rutishauser, U. Combined Phase-Rate Coding by Persistently Active Neurons as a Mechanism for Maintaining Multiple Items in Working Memory in Humans. Neuron 106, 256–264.e3 (2020).

51. Courellis, H. S., Valiante, T. A., Mamelak, A. N., Adolphs, R. & Rutishauser, U. Neural dynamics underlying minute-timescale persistent behavior in the human brain. 2024.07.16.603717 Preprint at 10.1101/2024.07.16.603717 (2024).

52. Mau, W. et al. The Same Hippocampal CA1 Population Simultaneously Codes Temporal Information over Multiple Timescales. Current Biology 28, 1499–1508.e4 (2018).

