## Supplementary material for "Time Cells in the Human Brain Support Working Memory Maintenance": suplemental figures

### Supplemental Figures

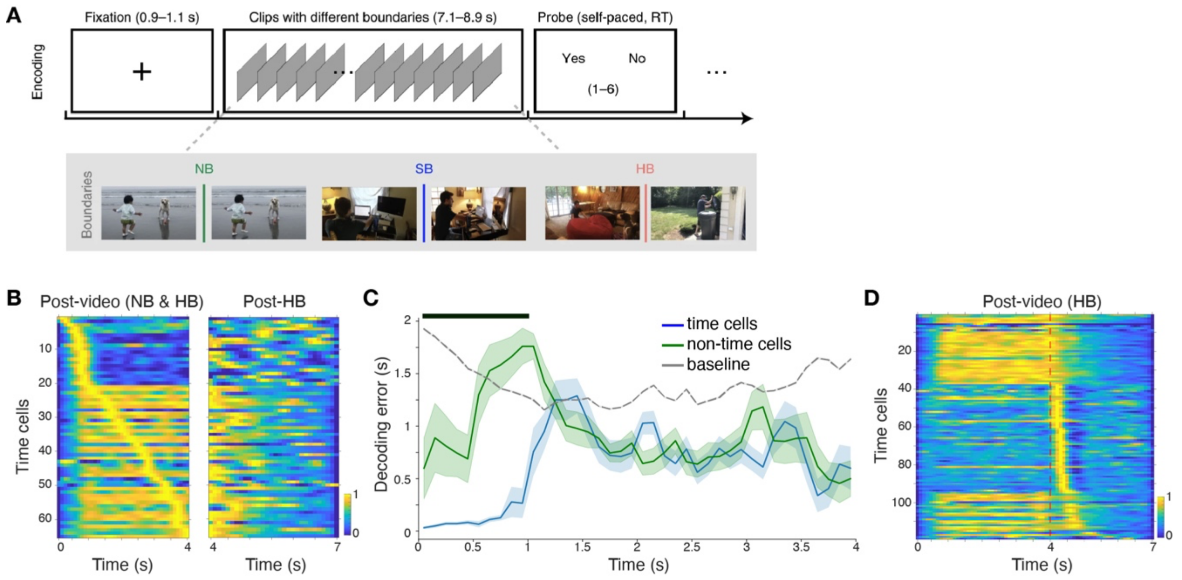


**Fig. S1 | Temporal coding in hippocampal and medial-temporal units in a non-working-memory task**. **A.** During each of 90 recording sessions, patients viewed videos containing either NB (no cut), or a SB (soft transition between scenes) or HB (abrupt narrative boundary) after the first 4 sec. **B.** Left: Average normalized firing rates of time cells, detected across the first 4 sec of boundary-free viewing in NB and HB videos, and ordered by peak latency in that window. Right: Average rates of the same neurons, sorted as before, after the boundary timepoint in HB segments. **C.** Bayesian time-decoding error for time cells (blue), non-time cells (green) and shuffled baseline (grey). Bar. Bins of significant difference between decoding errors of time cells and non-time cells ($p<0.05$). **D.** Averaged normalized firing rates of temporally modulated neurons detected across the entire duration of HB videos, sorted by peak latency post boundary (dashed line).

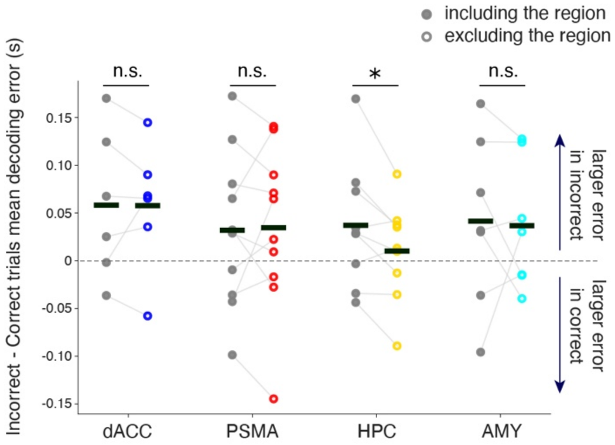


**Fig. S2 | Regional dropout analysis of temporal decoding during maintenance.** For each patient (included if ≥5 total time cells, ≥1 time cell in the dropout region, ≥1 time cell remaining after dropout, and ≥1 incorrect trial), we trained a Gaussian Naive Bayes decoder to estimate elapsed time during maintenance. For each region, decoding was performed once using all neurons and once with neurons from the dropout region removed, using identical train–test splits. Points show per-patient differences in mean decoding error between incorrect and correct trials (y-axis), with horizontal bars indicating region means (two-sided t-test; HPC: $N = 9$, $p = 0.035$; AMY: $N = 7$, $p = 0.869$; dACC: $N = 6$, $p = 0.998$; PSMA: $N = 10$, $p = 0.977$)

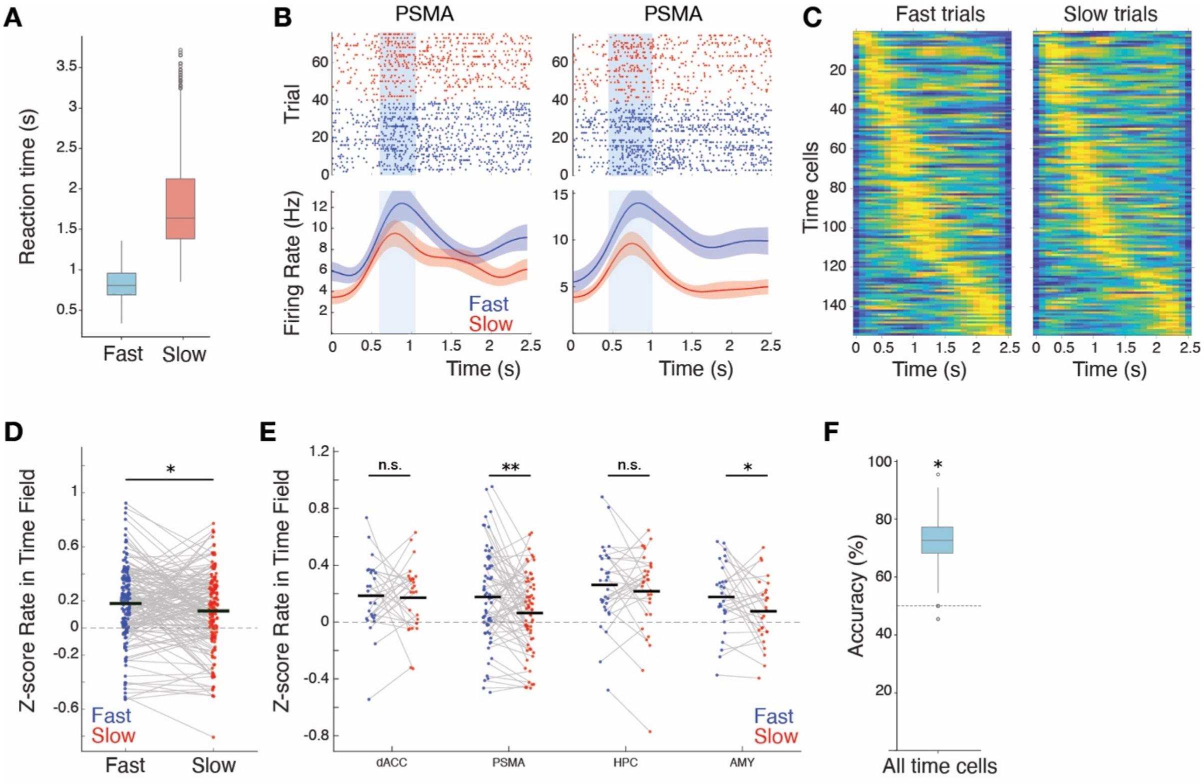


**Fig. S3 | Time cells predict probe reaction time in correct trials. A.** Reaction times in fast versus slow trials. **B.** Example units, plotted as in Fig 2 for slow vs fast response trials. **C.** Average normalized firing rates of time cells during correct trials with fast versus slow reaction times. Neurons are sorted based on time field over all trials. **D.** Within-field average z-scored firing rates in fast versus slow reaction time trials (paired t-test, $p = 0.0108$) over all neurons **E.** Within-field average z-scored firing rates in fast versus slow reaction-time trials (paired one-sided tests, Fast > Slow): $p_{dACC} = 0.410$; $p_{PSMA} = 0.002$; $p_{HPC} = 0.189$; $p_{AMY} = 0.025$**. F.** Decoding accuracy of reaction time (fast vs. slow) based on within-field firing rates using a Support Vector Machine classifier (empirical $p = 0.015$).

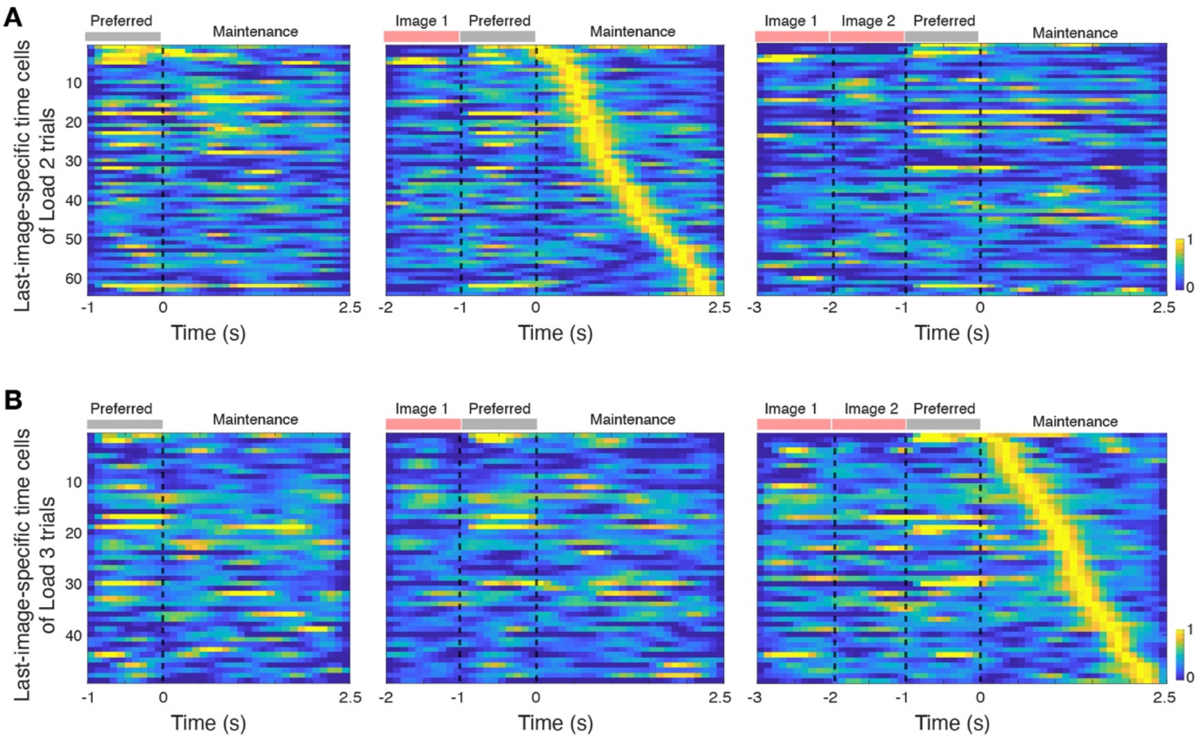


**Fig. S4 | Cross-load instability of "final-image"-selective time cells. A.** Average normalized firing rates of "time cells" detected based on their selective firing to a particular image when presented last in Load 2 trials (n = 64), sorted by their peak activation timepoint in these trials. The same cells, sorted the same way, are shown in Load 1 and 3 trials as well, when the same image was shown last. Firing rates are normalized by each neuron's peak rate during maintenance following the preferred image in Load 2. Dashed lines mark borders between images and maintenance onset. **B.** Same but for time cells selective to the last image in Load 3 trials, stacked by their field in those trials (n = 49).
